# Rhythmic modulation of dorsal hippocampus across distinct behavioral timescales during spatial set-shifting

**DOI:** 10.1101/2025.02.19.639177

**Authors:** Maeve Bottoms, Jesse T. Miles, Sheri J.Y. Mizumori

## Abstract

Previous work has shown frequency-specific modulation of dorsal hippocampus (dHPC) neural activity during simple behavioral tasks, suggesting shifts in neural population activity throughout different task phases and animal behaviors. Relatively little is known about task-relevant orchestrated shifts in theta, beta, and gamma rhythms across multiple behavioral timescales during a complex task that requires repeated adaptation of behavioral strategies based on changing reward contingencies. To address this gap in knowledge, we used a spatial set-shifting task to determine whether dHPC plays a specific role in strategy switching. The task requires rats to use two spatial strategies on an elevated plus maze: 1) alternating between East and West reward locations or 2) always going to the same reward location (*e*.*g*., only East or only West). Across specific timescales (session-based alignments, comparisons of trial types, within trial epochs), dHPC associated differentially with all three temporal categories. Across a session, we observed a decrease in theta and beta power before, and an increase in theta power after, the target strategy changed. Beta power was increased around the point at which rats learn the current rule. Comparing trial types, on trials before a rat learned the correct strategy, beta power increased. Within a single trial, after an incorrect (but not correct) choice, beta and gamma power increased while the rat returned to start a new trial. If gamma (but not beta) power was high during this return, the rat was more likely to make a correct choice on the next trial. On the other hand, low gamma power during the return was associated with incorrect trials. Rhythmic activity in dHPC, therefore, appears to track task demands, with the strength of each rhythmic frequency differentially associating with specific behaviors across three distinct timescales.

## Introduction

The dorsal hippocampus (dHPC) is traditionally known to be involved in memory (Eichenbaum, 1999), spatial navigation (Moser et al., 1995, Morris et al., 1982) and learning (Kim & Frank, 2009) The contribution of dHPC to these functions may reflect its role in the ability to flexibly switch between task strategies (*e*.*g*. Eschenko & Mizumori, 2007; Avigan et al., 2020). Such behavioral flexibility is typically measured in rodents via performance on a behavioral task that requires subjects to adaptively change response patterns, seen in reversal learning, spatial set-shifting, or attention set-shifting tasks (Flintoff et al., 2024, Riceberg et al., 2022, Miles et al., 2024, Kidder et al., 2021). In the case of reversal learning, the strategies are usually presented as opposing reward contingencies, without a change in the underlying rule (*e*.*g*. go west, then go east; maintains a strategy of going to a single reward location), while set shifting tasks require switching between more diverse strategies (*e*.*g*. a response strategy, then a place strategy). The structure of these tasks allows for analysis of neural activity preceding and following the strategy switch, providing a deeper understanding of the neural states that are supporting switches, and by proxy, flexibility (Miles et al., 2024), which is impaired in an array of psychiatric disorders, including addiction and depression (Maramis et al., 2021, Verdejo-Garcia et al., 2015). In the case of set-shifting tasks, place and response set shifting tasks provide a framework by which animals must switch between two distinct strategies, but there are contradicting reports of dHPC’s role in promoting egocentric or allocentric strategies (Packard & McGaugh, 1996, Gasser et al., 2020). Therefore, tasks requiring only the use of distinct and unique spatial strategies may offer a clearer insight into dHPC contributions to flexible strategy switching.

In dHPC, the different frequencies of local field potential (LFP) oscillations seem to reflect different aspects of behavior that may relate to flexible strategy switching. The theta rhythm (5-13 Hz) is often associated with attention and movement (Buzsaki, 2002) of the animal. The beta rhythm (15-30 Hz) is traditionally known for its involvement in sensory (Vanderwolf, 1992, Heale et al., 1994) and motor (Barone & Rossiter, 2021) behaviors, but is more recently thought to have an active role in cognition (Spitzer & Haegens, 2017) specifically involving the HPC (reviewed in Miles et al., 2023). Finally, the gamma rhythm (40-100 Hz) is frequently implicated in learning and memory processing (Colgin & Moser, 2010).

Evidence shows that dHPC LFP activity changes trial-to-trial, based on the demands of the task and the ongoing process of making decisions. Behavioral metrics such as vicarious trial and error (VTE) have neural correlates in dHPC both in single unit representations (Amemiya & Redish, 2016), and in predicting VTE behavior based on LFPs (Miles et al., 2021). On maze-based tasks, VTEs are articulated as a trajectory that initially trends towards one choice, then reverses direction to end up at a different location. VTEs are thought to be a deliberative behavior reflecting uncertainty with regard to an upcoming choice (Redish, 2016). Recent work shows that VTEs are likely to precede correct choices in a set-shifting task, with the presence of VTE behavior correlating with an increase in rhythmic power in the beta band of the medial prefrontal cortex (mPFC) (Miles et al., 2024). Similarly, inactivation studies in mPFC provide evidence that the disruption of mPFC results in dysregulation of VTE behavior (Schmidt et al., 2019, Kidder et al., 2021, Kidder et al., 2024). It appears that both dHPC and mPFC have supporting evidence for their involvement in producing or regulating VTE behavior, though it is not well understood how dHPC in particular contributes to VTE behaviors during complex set-shifting tasks requiring the use of different spatial strategies. In addition, studies report that task-related theta interactions between mPFC and dHPC are involved in choice accuracy (Stout et al., 2024, Jones & Wilson, 2005, Hyman et al., 2010), though little is known about whether activity in dHPC alone is enough to predict accuracy.

Recent advances in complex behavioral task quantification have led to the ability to identify a learning point during set-shifting tasks as rodents actively learn the correct strategy, allowing the separation of trials into two distinct categories of pre-learning (exploration), and post-learning (exploitation) (Maggi et al., 2024, Miles et al., 2024). Previously, measurement of exploratory behavior was often restricted to novel object recognition tasks (Lueptow, 2017), rather than reward-based tasks. While HPC neural network activity was found during these exploratory phases of object recognition (Franca et al., 2014, Iwasaki et al., 2021), our understanding of exploratory phase neural activity during complex reward-based maze task performance is not yet known at the level of precision presented in Miles et al., 2024.

Within a single trial, the LFPs of dHPC are known to show an increase in theta power during the decision epoch compared to the “control” epoch within trials (Schmidt et al., 2013, Jones & Wilson, 2005), but the relevancy of these within-trial epoch modulations on future behavior is not yet clear. Human and non-human primate literatures implicate the gamma band in learning and memory processing after presentation of a stimulus (Sederberg et al., 2007b, Jutras et al., 2009), as well as before a choice (Sederberg et al., 2007a) in recognition memory tasks, indicating that increases in power at different times during a trial could reflect differential memory processing. Since maze-based trials can be separated into distinct epochs before and after reward, the role of the rhythmic activity in task performance can be further dissected to see if positive or negative reward feedback is reflected in changed rhythmic power throughout other trial epochs.

This study aimed to clarify the roles of dHPC theta, beta, and gamma rhythms relative to different temporal scales of behavior during a spatial memory task that requires rats to adapt their strategy several times throughout the session. LFPs were recorded while rats performed a spatial set-shifting task that required repeated, uncued, switches between two spatial strategies. This design allowed us to examine LFP changes on different temporal scales of the task, including session-based alignment, comparison of trial types, and within-trial changes. This framework enabled an understanding of dHPC contributions across different levels of task granularity. We hypothesized that theta would correspond to session-based alignment metrics, while the beta and gamma frequencies would reflect learning and cognition processes relevant to individual trials or behaviors, manifesting as associations between power and reward feedback. Results from these investigations show that, when examining session-based alignment, rhythmic power appears to track task demands. Theta and beta power both decreased preceding an update in task context, while theta increased after this update. Beta power increased around the time rats learned the current rule. A comparison of trial types revealed higher beta power while rats executed a choice prior to learning compared to post-learning trials. Within a single trial, beta and gamma power increased after an incorrect choice as rats returned to the start location. This modulation of gamma power specifically during the return epoch was associated with choice accuracy on the subsequent trial, and this relationship scaled with the strength or weakness of gamma power. In sum, we show that dHPC rhythmic power varies on three distinct temporal scales and reflects the intricacies of reward processing and real-time strategy updating.

## Methods

### Subjects, apparatus, training protocol, and behavioral task

Long-Evans rats (n=5, 3 female; Charles River Laboratories) were food restricted to 85% of ad libitum body weight with free access to water. Rats were housed in a temperature and humidity-controlled facility with a 12 hr light-dark cycle. Behavioral testing occurred during the light cycle, and training took place when rats were 4-12 months old. All care and procedures were performed in compliance with the University of Washington Institutional Animal Care and Use Committee under National Institute of Health guidelines.

The apparatus, training protocol, and behavioral task were much the same as Miles et al., 2024. In short, the task was performed on an elevated plus maze (black plexiglass arms, 58 cm long x 5.5 cm wide, 80 cm from the floor) with remotely moveable arms and reward ports controlled via a custom LabView 2016 software (National Instruments). The software controlled movement of the arms and dispensed rewards based on the recorded position of the rat measured by a SONY USB camera (Sony Corporation) capturing frames at ∼35 Hz. After arriving in our facility and being acclimated to handling and food restriction, rats were placed on the maze with scattered 45 mg sucrose pellets (TestDiet) to allow them to habituate to the room, maze, and maze movements. Rats were allowed to explore the maze for 10-15 minutes daily for at least 3 days before moving on to pre-training.

During pre-training, rats initially trained according to a forced-choice paradigm to encourage the acquisition of the trial structure and task rules. After leaving a pseudo-randomly chosen start arm (“North” or “South”), rats were forced to navigate to either the “East” or “West” reward arms to receive a guaranteed sucrose pellet reward, according to the designated strategy in each block. After receiving reward, rats returned to the next pseudo-randomly available start arm. Each pre-training session consisted of one block of each strategy type, each block of which included 5 forced-choice trials. That is, choices either alternated between reward arms, or remained the same (East or West), regardless of start location. The order of presented strategies was randomized for each session. As rats successfully completed all trials in a session, the block length of each strategy increased in subsequent sessions, until rats were completing 15 trials of each strategy type for a combined 45 trials in 45 minutes.

After completing the above pre-training (approximately two weeks of training), rats entered into a free-choice version of the task. In this paradigm, rats were presented with a choice between East and West reward arms and learned to recognize uncued changes in reward contingencies in order to complete the task. Each session consisted of 4 blocks: two alternation blocks, one place East block, and one place West block. Block sequences were generated such that the two alternation blocks within a session could not be adjacent. For example, ‘Alt-W-E-Alt’ and ‘W-Alt-E-Alt’ were possible, but ‘W-Alt-Alt-E’ or ‘W-E-Alt-Alt’ sequences were not. In order to progress to the next block, rats were required to reach a choice accuracy criterion in a sliding window of 10-15 trials; the experimenter set and adjusted this window as rats learned the task. Rats began training with a smaller sliding window (10-12 trials) but were required to reach an accuracy criterion of 80%. As rats became proficient in the task, the window was increased, thereby increasing the difficulty, until rats made 12/15 correct choices in order to progress to the next block, referred to here as the block switch. Sessions were capped at 150 trials or one hour for training. If rats successfully completed all four blocks within 150 trials, the program automatically shut down and the session ended.

### Electrophysiological implant and recordings

Custom-built tetrode (NiCr, Allemia) microdrives were constructed with Intan headstages and contained 32 channels localized to dHPC. To prevent electrical damage, the microdrives were glued into plastic tubes lined with copper tape. One ground wire was secured to the copper-taped tube using silver epoxy, and the other ground wire was soldered onto a screw to be implanted over the cerebellum. After completing four blocks in a session on two different days, rats underwent a surgery to implant the custom-built microdrive that contained tetrodes aimed at dHPC (AP = -4.5mm, ML = 1.8mm, DV = -1.7mm). A subset of rats (n=4) had an additional microdrive that contained channels localized to the prefrontal cortex (those data are not described here). After full recovery from surgery, rats were retrained on the task before recording sessions began. Recordings were collected similarly to Kidder et al., 2021 and Miles et al., 2024 via an Open-Ephys acquisition system acquiring data at 30 kHz. Tetrodes were lowered after each session until channels exhibited large amplitude theta rhythms so that all recordings were localized to a similar area right below the pyramidal layer of dHPC CA1. A total of 17 sessions were analyzed, with each rat contributing between two and four sessions.

### Tracking and VTE classification

An LED was placed on the animal’s head during electrophysiological recording sessions. Sessions were recorded with a Blackfly camera (Teledyne Vision Solutions; ∼60Hz) for position tracking; the video was later processed using a custom MATLAB script to decode pixels into centimeters and center the maze on a Cartesian plane for position tracking. A .csv file containing the position of the rat during each frame was outputted and was used to classify VTE behaviors in the same way as previously described in Kidder et al., 2024 and Miles et al., 2024. In short, the position data were aligned to the choice point, truncated to have the same start and end coordinates, and then interpolated (or linearly subsampled, if required) to have equal numbers of points for analysis. Trajectories were then projected into principal component space and clustered according to the trajectory representations using hierarchical agglomerative clustering. Clusters were separated using distance-based dendrograms to result in two clusters of VTE and non-VTE trajectories. Misidentified trajectories were corrected based on x-position coordinates and the *z-ln(idphi)* measure (Papale et al., 2012, Papale et al., 2016). Also, several lab members visually inspected the VTE clustering output and it was determined that the automated clustering method fell between the 80-90% accuracy threshold (relative to manual labeling) expected.

### Behavioral and electrophysiological analysis

For each session, strategy likelihoods were generated (similar to Maggi et al., 2024) for each strategy (alternate, go East, and go West) using recency weighted Bayesian inferencing on the success or failure of each strategy type for each trial. This generated a metric for the likelihood that an animal would use a given strategy on each trial. Within each block, the learning point was classified as the trial at which the correct strategy became the most likely strategy. As a new block started, trials for which the previously correct strategy likelihood continued to increase comprised the perseverative phase. Trials between the perseverative phase and the learning point were classified as exploration trials, while any trials after the learning point and before the next block switch were classified as the exploitation phase (*Figure 1b*).

**Figure 1.**
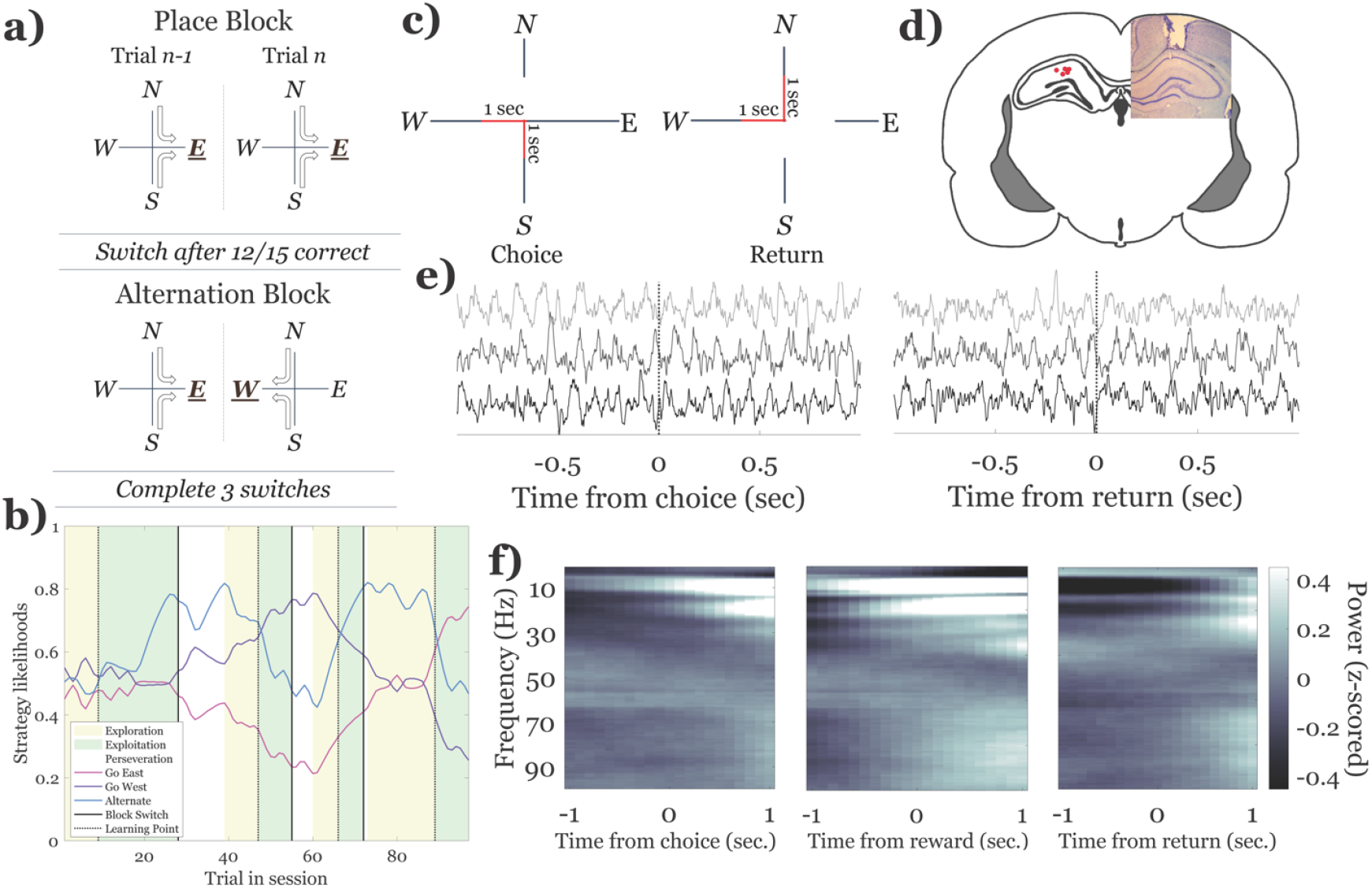
Description of the task and electrophysiology. a) Diagram of task rules. Rats start on a randomly selected block of trials, either place or alternation. The place strategy requires animals to go to the same reward arm each trial (either East or West, regardless of start location). The alternation strategy requires animals to alternate between reward arms (East, then West). Start arms (North and South) were pseudo-randomly chosen after every trial, ensuring rats used a spatial strategy. A block switch occurs when rats achieve 12/15 trials correct. A session consists of four blocks with three block switches. b) Strategy likelihoods and phases across a single session. One session from one rat was plotted as a demonstration of the calculated likelihoods and phases of the task. See Methods for more details on calculations. c) Depiction of the choice and return epochs. During the choice epoch, rats ran from a start arm (North or South) to a reward arm (East or West). During the return epoch, rats received reward feedback and ran back to start a new trial at a new start arm. Both epochs encompassed 1 second before and after crossing the platform. d) Histological verification of tetrodes. Recording sites were localized to the fissure of the dorsal hippocampus. e) Electrophysiological data were aligned to the maze center for both choice and return epochs. Example LFP traces aligned to the choice and return epochs. f) Averaged spectrograms from all trials aligned to choice, reward encounters, and return.

Electrophysiological data were separated into two non-overlapping windows: choice and return. The choice window was determined by first locating the time point where the animal was closest to the center platform during their choice trajectory; the return window was centered on the time point where the animal was closest to the center platform during their return trajectory. A third time window, reward, was plotted to demonstrate high theta power during periods of stillness. Four seconds of data before and after the center crossing were used to calculate a time-frequency spectrogram using Chronux’s mtspecgramc function (Boikil et al., 2010; two second window, 100 msec overlap, seven tapers, and a bandpass window of 0.5-100 Hz). This created a time-frequency matrix for each choice and return trajectory. Trials with excessive noise were identified by visual inspection and discarded based on input from several experimenters (0.015% of trials for choice, 0.042% of trials for return). Power values were converted to decibels, then Z-scored across each individual time window to normalize the data. Data were then reduced to +/-one second from the choice or return point, both of which were considered to be the central-most position on the center platform of the maze, and averaged for each trial to produce one value for choice and return.

Data shown in *Figures 2, 3 & 4* were sampled according to a hierarchical bootstrap method presented by Saravanan et al., 2020, and implemented in the same way as Miles et al., 2024. The number of iterations for each bootstrap distribution was set at 1,000, while the number of samples and sessions for each iteration was set at or below the lowest value observed across all sessions and rats. For the bootstraps presented in *Figure 2*, samples were taken at the rat and block levels, with replacement. An entire sequence of 21 trials was taken as a single sample at the block level to accurately represent the temporal dynamics of each rhythm aligned to a specific time point during the session. For one iteration, subjects were randomly sampled up to the total number of subjects in the dataset (five). From each subject, six learning points and eight block switches (equivalent to two complete sessions) were sampled, then used to extract the corresponding electrophysiological data. This process was repeated for 1,000 iterations. Plots presented in *Figures 3 & 4* were sampled at the rat, session, and trial level, with 40 samples taken from each session for *Figures 3* and *4a*, 15 samples for *Figure 4b*, and ten samples for *Figures 4c-d*. Sample numbers varied based on the lowest number of trials of each type present in all blocks, to ensure no repeat sampling during iterations. These plots were created by comparing the rhythm power (*Figure 3&4a*) or accuracy (*Figure 4b,c,d*) on different trial types. For example, for a histogram depicting the association between rhythm power and accuracy (*Figure 4a*), correct trials were designated to be “condition” trials, while incorrect trials were “comparison” trials. This created within-subject pairs of trials for each corresponding response variable (in this case, power). The means of each response variable were subtracted from each other during each iteration, creating a distribution, with 0 reflecting no difference between the condition and comparison trial types. This method ensured equal contributions from each rat, regardless of the number of sessions they successfully completed. To create the plots shown in *Figure 4b*, the same bootstrap analysis was used to compare trials with low and high power for each rhythm. Trials were separated based on percentiles of rhythm power, ensuring that there were the same number of low and high trials across the dataset to draw from (*e*.*g*. 50/50, 40/60, 25/75). This created a group of trials that was within the lower percentile, and another group within the higher percentile, used as the condition and comparison filter for the bootstrap. Choice accuracy on the next trial was used as the response variable instead of power, which was used in *Figure 3, 4a*. This created plots demonstrating how choice accuracy varied after trials with low or high power during the return epoch.

**Figure 2.**
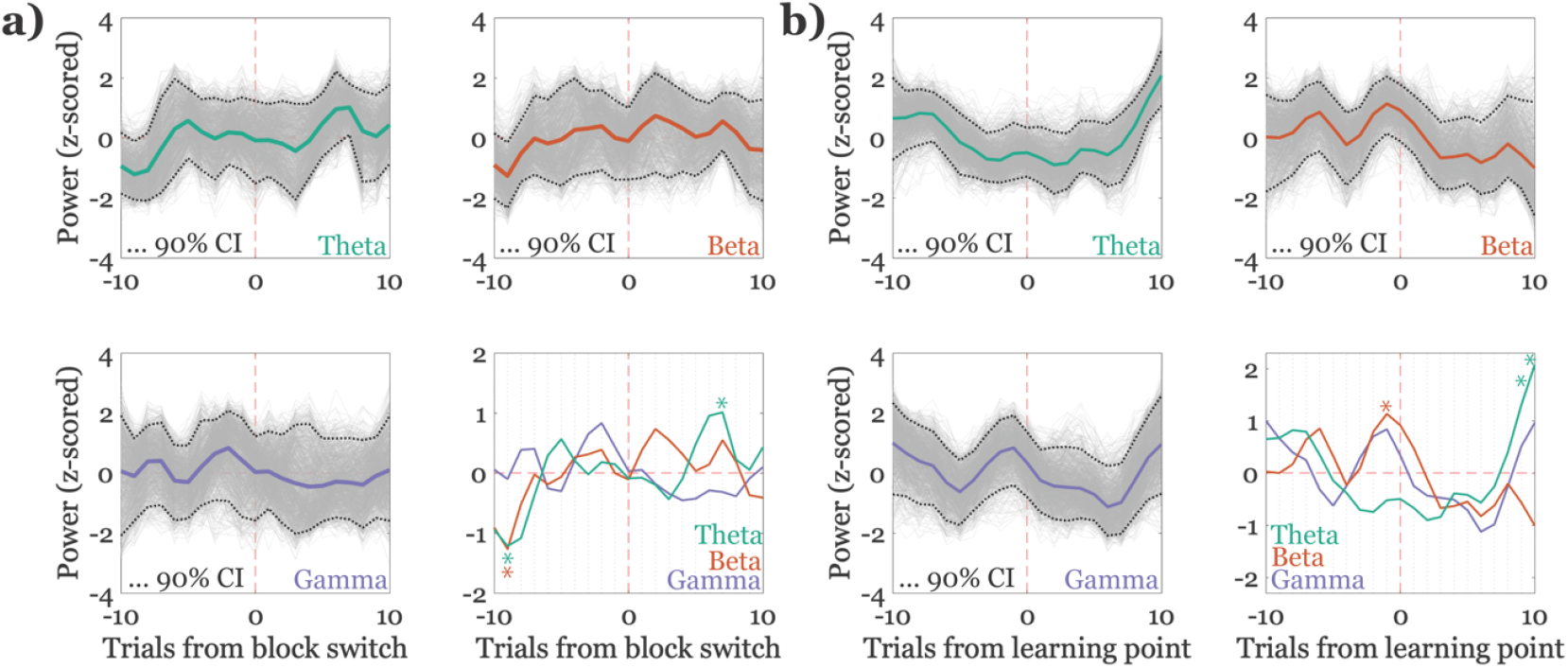
dHPC activity during choices changes with respect to task structure and contingency learning. Asterisks represent trials in which 95% of iterations fell above or below zero and the standard deviation was >1.5. a) Rhythmic power aligned to the block switch. Power values from ten trials before and after an uncued strategy change (block switch) were plotted using a hierarchical bootstrapping method. In the upper two boxes as well as the lower left box, 1,000 iterations of the bootstrap are shown in gray for each frequency band, with the average of the iterations shown in color. 90% confidence intervals (CI) are shown as the dotted line. In the lower right box, the sample averages for each frequency band are plotted together. b) Rhythmic power aligned to the learning point. The learning point was defined as the trial at which the correct strategy became the most likely for the animal to use (see Methods). Ten trials before and after the learning point were plotted for each of theta, beta, and gamma. The lower right panel compares the three frequency bands.

**Figure 3.**
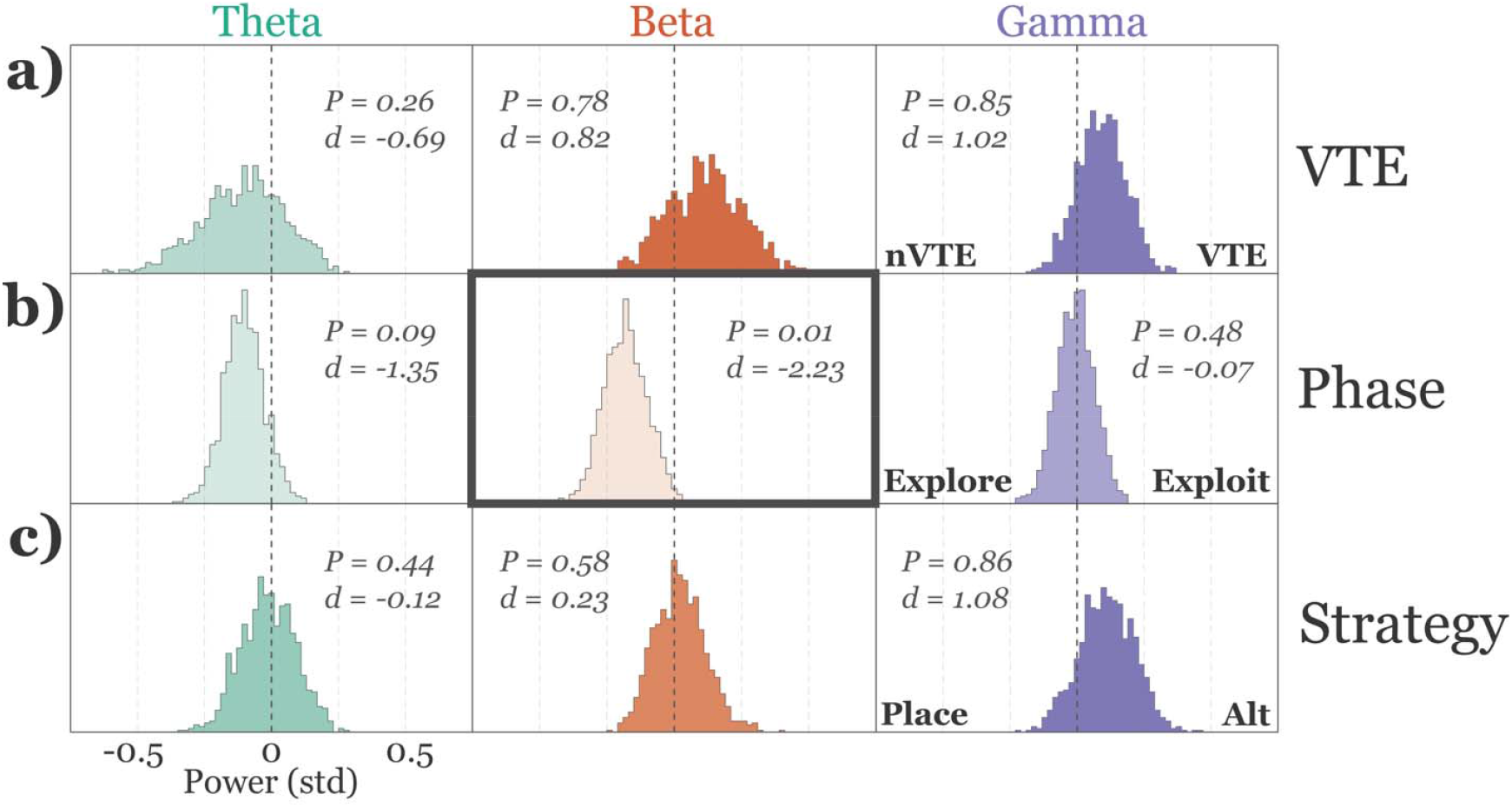
Choice epoch dHPC power comparison across different trial types. Each box depicts an association between a particular frequency band and a trial type. Histograms were plotted based on a hierarchical bootstrap algorithm that subtracted the power values from paired trial type comparisons (*e*.*g*. theta power during a non-VTE (nVTE) trial was subtracted from theta power during a VTE trial). See Methods for more details. P values were calculated as the proportion of values that fell above or below zero, while d values were calculated as the standard deviation. A P value of >0.95 or <0.05 combined with a d value of magnitude >1.5 was considered to be statistically significant. Significance is illustrated by the bold-faced box outline. a) dHPC power does not strongly associate with VTE behaviors. VTE trials do, however, tend to have higher gamma power compared to non-VTE trials. b) dHPC beta power associates with exploratory trials. Theta power shows a similar trend, but less so than beta. c) dHPC power does not strongly associate with strategy type. Alternation blocks appear to moderately associate with high gamma power but does not meet the threshold for a significant association.

**Figure 4.**
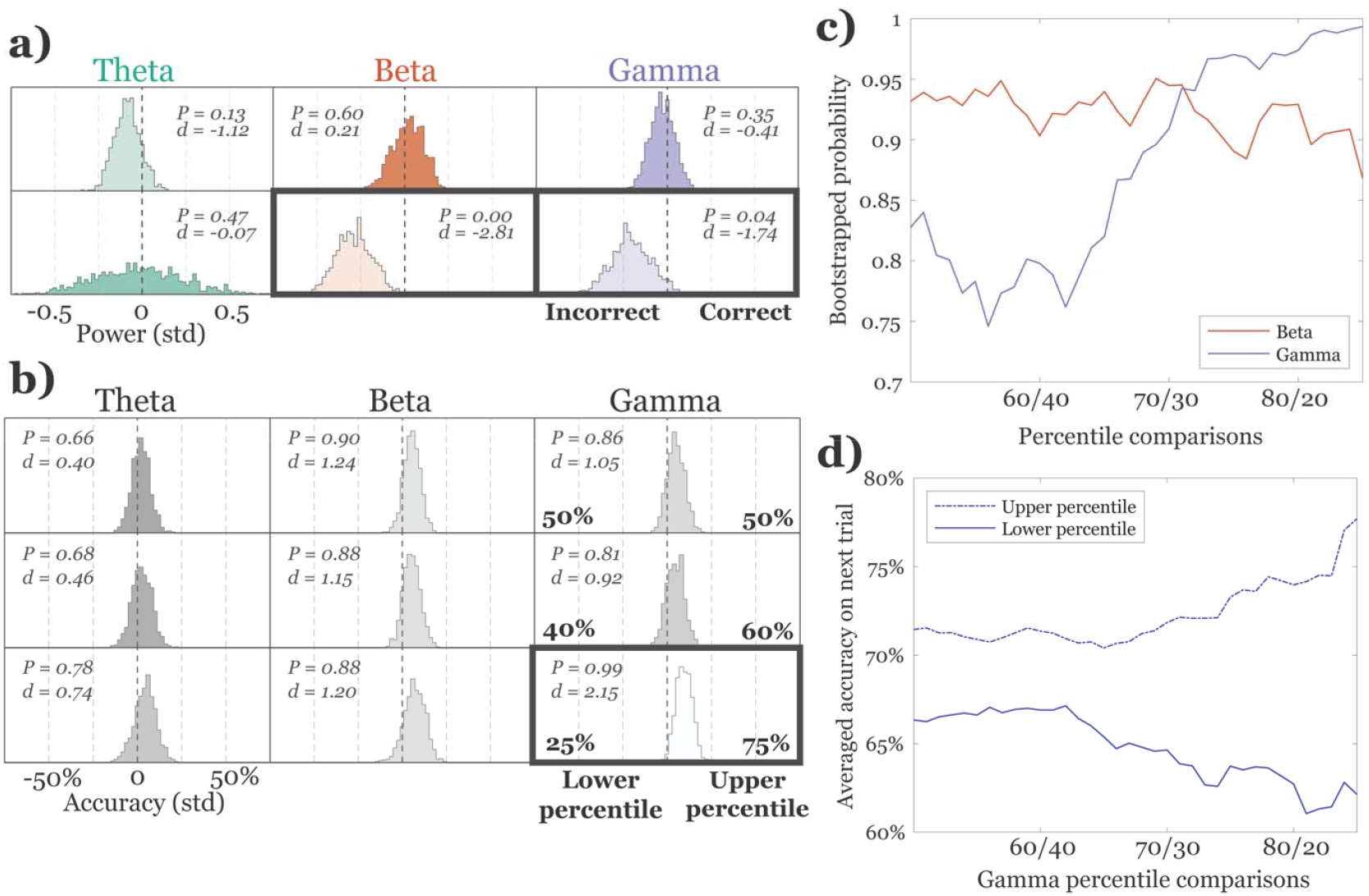
Using the analysis described for *Figure 3*, associations between LFP frequency bands and choice accuracy are shown. a) Beta and gamma power are higher during the return epoch after incorrect choices, but not during the choice epoch. b) Gamma, but not beta, power in the 75th percentile significantly associates with greater accuracy on the subsequent trial. A similar bootstrap to *Figure 3* was run, showing no significant associations between different percentile ranges, except for in the 25th/75th percentile gamma comparison. c) The likelihood of higher accuracy increases as the threshold for high gamma power, but not beta power, increases. The y-axis represents the probability of accuracy values falling above 0, indicating greater accuracy on the higher percentile threshold trials. The x-axis represents the percentile comparisons. The gamma distribution shows a clear increase around x=13, while the beta distribution appears to be random and noisy. d) Accuracy on a subsequent trial scales with thresholds for high and low power gamma. Actual averaged accuracy values were plotted from the lower and upper percentile thresholds, showing that as the definition of low and high become more stringent, the accuracy of the next trial scales in the same direction as the corresponding power percentile.

### Statistical analysis

For all bootstrapped plots (*Figures 2, 3 & 4*), P-values were computed as the proportion of samples that fell above or below zero, while d values were computed as the mean of the sample distribution divided by the standard deviation of the same distribution. The null hypothesis in these cases was taken to be P=0.50. Since an alpha of 0.10 was used on two-sided distribution plots, a distribution with 95% of samples falling above *or* below zero, combined with d>1.5 was considered to be statistically significant. P values were calculated without the use of traditional hypothesis testing. Therefore, a lower P of 0.80 (i.e., 80% of values fall above or below zero) still provides some support to reject the null hypothesis.

## Results

To evaluate the role of dHPC in complex strategy-switching behavior, we recorded LFPs (*Figures 1d-f*) during a spatial set-shifting task performed on an elevated plus maze that required rats to use both place and alternation strategies (Miles et al., 2024). Sessions started randomly with one strategy, and after reaching 12/15 trials correct in a sliding window, an uncued rule change occurred, and rats adapted to the new required strategy (*Figure 1a*). Following the logic of Kidder et al., 2024, data were separated into two distinct, non-overlapping epochs. The choice epoch was segmented by finding the time when rats crossed the center platform *en route* to a goal, representing the time of their choice; the return epoch was aligned to the same physical location in space, but represented times when rats crossed the center platform after receiving reward feedback while returning to a start arm to begin a new trial (*Figure 1c*). Data were windowed to include one second before and after crossing the center platform (*Figure 1e*). Averaged time-frequency plots across all trials for both epochs can be seen in *Figure 1f*.

Three neural frequency ranges of interest were extracted from the data: theta (5-13Hz), beta (15-30Hz), and gamma (40-100Hz). Since each of these rhythms have been implicated in learning and memory tasks in the hippocampus (Buzsaki, 2002, Miles et al., 2023, Colgin & Moser, 2010), we examined the activity of these specific frequency bands in the context of the overarching task structure: Trials were aligned to either the block switch or learning point, and the power for each rhythm was hierarchically bootstrapped during specified time windows (see Methods). *Figure 2a* shows data aligned to the block switch. For each rhythm, the iterations of the bootstrap algorithm were plotted with a 90% confidence interval surrounding them.

Aligning all choice epoch rhythmic activity to block switches revealed a significant decrease in theta and beta power 9 trials before the block switch (*Figure 2a*, theta trial -9, P=0.03, d=-1.96, beta trial -9, P=0.04, d=-1.85), while theta power significantly increased on trial 7 after the block switch (*Figure 2a*, theta trial 7, P=0.97, d=2.03). Relative to the learning point, beta power significantly increased 1 trial before the learning point (*Figure 2b*, beta trial -1, P=0.97, d=1.96) while theta power increased 9 and 10 trials after the learning point (*Figure 2b*, theta trial 9, P=0.99, d=2.74, theta trial 10, P=1.00, d=3.59). These general trends suggest that during choice behavior, theta power increases after both the block switch and learning point. The beta rhythm clearly exhibits a brief elevation preceding the learning point. Gamma power during choices was not clearly associated with either changes in strategies or learning.

To compare different trial types in the context of each frequency band during the choice epoch, a similar hierarchical bootstrapping algorithm that randomly and equally sampled paired trial types at the level of rat, session, and trial was used. The paired trial types selected were VTE/non-VTE trials, exploration/exploitation trials, and trials during alternation/place blocks. For each comparison, the power value of the condition trial (VTE, exploitation, and alternation trial types) was subtracted by the value of its respective comparison trial (non-VTE, exploration, and place trial types). The resulting value was either positive, indicating higher power on the condition trials, or negative, indicating higher power on comparison trials. The bootstrap algorithm was run for 1,000 iterations, and the results were plotted as histograms, showing the distribution of comparisons. If a histogram falls on the right side of the bold vertical dashed line, it denotes higher power during the condition trial, while a histogram on the left side indicates higher power during the comparison trial. The d and P values were calculated based on the standard deviation and proportion of trials falling on each side of the plot. A histogram with a P value of less than 0.05 or greater than 0.95, combined with a d of magnitude greater than 1.5 was considered to be significant evidence for an association. Based on these criteria, there was no association between dHPC activity and VTE behavior (*Figure 3a*), though there was a trend towards increased gamma power on VTE trials (*Figure 3a*, gamma, P=0.85, d=1.02). When examining exploration and exploitation trials, there was a significant association between exploratory trials in the beta frequency band (*Figure 3b*, beta, P=0.01, d=-2.23), when compared to exploitation trials. Theta power appears to demonstrate a similar pattern, though with less consistency (*Figure 3b*, theta, P=0.09, d=-1.35). This indicates higher theta and beta power before the learning point, consistent with the alignment plots shown in *Figure 2*. Finally, we compared strategy types and found no strong associations, though there was a trend toward higher gamma power during alternation blocks (*Figure 3c*, gamma, P=0.86, d=1.08). Taken together, these data show that, while dHPC rhythmic activity associates with different trial types, the strongest association occurred during exploration periods, *i*.*e*., before the target strategy is learned.

The rat’s goal during the spatial set shifting task is to maximize rewards. To do so, rats must incorporate information about choice outcomes to update their strategy. By plotting the same histograms seen in *Figure 3*, but this time comparing correct and incorrect choices, there was no association between choice accuracy and rhythmic dHPC activity while rats carried out their choice. During the return epoch, however, after rats received choice outcome information and were preparing to begin a new trial, high beta and gamma power was significantly associated with incorrect trials (*Figure 4a*, beta, P=0.00, d=-2.81, gamma, P=0.04, d=-1.74). To determine whether this increase in power after an incorrect choice was predictive of future choices, comparisons were run by extracting choice outcome on the next trial after high or low beta or gamma power during the return epoch. By extracting different percentile ranges across all trials (*e*.*g*., comparing the upper 50th to the lower 50th, the 60th to the 40th, and the 75th to the 25th), we found that using a more stringent threshold for defining “high power” revealed a significant relationship (P=0.99, d=2.15) between high gamma power and choice accuracy on the next trial compared to its “low power” counterpart (*Figure 4b*). Notably, though beta power was also higher after incorrect choices on the return epoch (*Figure 4a*), it was not predictive of accurate choices on the next trial. Therefore, the association of high power during return epochs after an incorrect choice with a correct future choice was specific to gamma.

To further characterize the association between high gamma during a return epoch and subsequent choice accuracy, the impact of varying the threshold for defining high power was tested, ranging from the 50th to 85th percentiles of the beta and gamma range (*Figure 4c*). By plotting all comparisons, we found that a difference in accuracy between high and low gamma power reaches, and maintains, significance (P>0.95, d>1.5) starting when data are split between the 74th/26th percentiles,. This effect remained significant through the 85th/15th percentile comparison, at which point the comparisons no longer retained sufficient trials for accurate calculations. The accuracy plots in *Figure 4d* show both an increase in accuracy after high gamma power returns, and a decrease in accuracy after low gamma power returns. This observation could be enhanced by the increased likelihood that rats will make a correct choice after an incorrect choice, but the percentile regression does not take into account information about the previous choice, and the same effect is not seen in the beta frequency range (*Figure 4c*), even though beta power is similarly increased during the return epoch after an incorrect choice. These data indicate that strong gamma power during the return epoch is important for subsequent choice improvement.

## Discussion

We found that different frequencies of rhythmic dHPC activity fluctuated with respect to different time scales during a spatial set-shifting task. Theta rhythmic activity changed as reward contingencies updated during session-based alignments, such as block switches and the learning points. Beta activity, on the other hand, was elevated as rats learned to update behavior in response to reward-contingency changes, as well as in response to single-trial reward feedback. Like beta, within-trial gamma rhythmic activity was elevated after errors. Unlike beta, the strength of gamma activity after rewards was associated with the likelihood of a correct choice on the next trial – lower gamma power during the return epoch was associated with more incorrect choices on the next trial, while higher gamma power led to an increase in accuracy on the next trial (*Figure 4*). These results indicate that, during a spatial set shifting task, dHPC activity associates with behavior across three distinct timescales (session-based alignments, seen in *Figure 2*, comparisons of trial types, seen in *Figure 3*, and within single trials, seen in *Figure 4*). Comparing these timescales allows insight into the role of dHPC during reward processing and strategy updating: the gamma rhythm seems involved in trial-to-trial learning, with the strength of power during the return epoch directly associating with subsequent accuracy. The beta rhythm appears to be associated with session-wide general strategy learning and application, as demonstrated by higher beta power during exploration trials and around the learning point.

Importantly, beta operates on all three timescales: session-based alignment around the learning point, during exploitative (not exploratory) trial types, and within a single trial after receiving incorrect reward feedback. It is worth noting that during task performance, we found no correlation between rhythmic power and velocity, indicating that the results described here could not be accounted for by the running speed of the animal per se. For example, *Figure 1f* shows high theta when animals were moving or when they remained still around reward delivery i.e.a time when animals were consuming or waiting for rewards to be delivered. Recent analyses of theta sequences during reward-related immobility showed that these sequences represented other locations in space, particularly the trajectory of the upcoming trial (Wang et al., 2025).

### Session-based alignment activity of the dorsal hippocampus

The structure of the spatial set-shifting task allows for a detailed observation of field potential responses as rats flexibly adapt to a sudden change in task context, or as rats are nearing the point of learning the current rule. Several studies show that mPFC responds to changing task contingencies (Karlsson et al., 2012, Powell & Redish, 2016, Malagon-Vina et al., 2018), but it was proposed that these changes in activity reflect errors or task performance instead of specific strategies a rat must use. Hasz & Redish 2020 showed that dHPC single-unit ensembles encode current task strategies, as well as active behavioral shifts, similar to our finding of neural changes relative to the learning point. Similarly, Ding et al., 2025 found that the single-unit dynamics of HPC tracked strategy switches throughout a session. Combining mPFC and dHPC, Mugan et al., 2024 found that activity in mPFC was most closely associated with dHPC activity after a rule switch. Around the learning point, we observed an increase in beta power, consistent with the observations of Rangel et al., 2015, who found an increase in beta amplitude, along with a decrease in theta amplitude, during associative learning tasks. Our task shows a similar link to learning, but in the context of rule switches. The design of our task allows for more detailed insights, tailored specifically to the block switch and learning point. We observed a decrease in theta and beta power before the block switch, and an increase in theta power after the block switch. It is possible that the beta rhythm contributes most to learning new task rules, culminating in a peak in power around the learning point, while the theta rhythm seems to respond most strongly to the block switch, when the rats are attending to updates in task context.

### The contribution of the dorsal hippocampus depends on trial types

We did not find differences in dHPC rhythms according to strategy types, which may be expected given that both strategies require spatial working memory. Several studies have, however, implicated mPFC in spatial alternation and place tasks (Sapiurka et al., 2017, Kidder et al., 2021, Kidder et al., 2024), and mPFC/dHPC interactions during spatial working memory have been well documented (Eichenbaum, 2017, Stout et al., 2024), indicating that the rhythmic activity of dHPC alone may not be enough to reflect specific strategy use. Further, we did not separate analyses based on trials after the learning point, but rather all trials in a block, which may have precluded strategy identification. Along these same lines, and contrary to prior reports (Amemiya & Redish, 2016, Miles et al., 2021), we saw no difference in rhythmic dHPC activity between VTE and non-VTE trials. Part of this discrepancy could be related to the prevalence of non-deliberative VTEs in this task (Miles et al., 2024), which we could not separate from the typical deliberative VTEs since some rats did not have enough VTEs in each category to make meaningful conclusions. Interestingly, work examining inactivation of mPFC in a different complex spatial working memory task (spatial delayed alternation) showed that, while mPFC optogenetic disruption did not decrease overall VTE rates, it did disrupt the timing of VTE with respect to reward contingency changes (Kidder et al., 2024). When using the same spatial set shifting task as the one from this study, mPFC beta power differed between VTE and non-VTE trials (Miles et al., 2024), again suggesting that mPFC may mediate VTE behavior in tasks requiring continual updating of spatial behavior.

In the present study, beta power increases were strongly associated with exploratory trials, which occurred before the learning point. Although the beta rhythm is traditionally linked to sensory systems (Boeijinga & DeSilva, 1989), several reports have described higher beta activity in the HPC during novelty or exploratory activity (Berke et al., 2008, Franca et al., 2014, Iwasaki et al., 2021). More specifically, HPC beta has also been implicated in linking sensory learning and memory-guided decision-making (reviewed in Miles et al., 2023). Here, we expand on previous findings by showing an observable increase in beta power as rats explore potential future strategies through trial-and-error decision-making. This is in line with the idea that in addition to its involvement in sensory exploration (Symanski et al., 2022, Martin et al., 2007), beta may also contribute to attention during the exploratory phase of a reward-based task in service of spatial memory processes and learning.

### Different roles for beta and gamma during working memory processes

In addition to playing a role during exploration, beta also operates on a shorter behavioral timescale. We show here that both gamma and beta rhythmic power increased after rats made an error and were returning to a start location to initiate a new trial. Based on the model described by Kidder et al., 2024, the return epoch is thought to represent an encoding period, while the choice epoch is thought to represent a retrieval period. Because both beta and gamma were elevated during the return epoch, both frequencies are likely associated with encoding the salience of reward feedback (*Figure 4a*). Beta, as discussed previously, seems to be involved in learning processes since an increase in exploratory beta activity links to better memory performance during object recognition recall (Iwasaki et al., 2021). Similarly, Symanski et al., 2022 found that during an odor association task, dHPC and mPFC units phase lock to the beta rhythm, especially prior to correct choices. Here, however, we show higher beta power after *in*accurate decisions, indicating that beta may be acting as a type of prediction error signaling mechanism in dHPC (Mizumori et al., 1999, Mizumori et al., 2000). This prediction error signal may operate in tandem with, or arise from, dHPC’s interactions with other structures, given recent work that has shown beta’s involvement with reward processing in associative memory tasks (Lansink et al., 2016, Igarashi et al., 2014). Beta, however, had no discernable association with accuracy on the next trial, indicating that this prediction error signal may act more as a modulator of memory by enabling a neural state that facilitates gamma encoding of information contained in the return epoch. This facilitation mechanism could help store memories of task-salient events in a way that is more accessible during subsequent retrieval.

Previous research on humans and non-human primates has shown that gamma power in the hippocampus is associated with both the execution of choice (Sederberg et al., 2007b), as well as the encoding of task-relevant information (Sederberg et al., 2007a, Jutras et al., 2009). Gamma is also linked to location and accuracy prediction in animal behavior. For instance, recent work showed that the individual features of gamma rhythms could be used to predict the location of rats on a simple radial arm single-goal maze task (Douchamps et al., 2024), suggesting that gamma has a role not only for learning, but in decoding and predicting animal behavior. Similarly, Dvorak et al., 2018 showed an increase in 30 - 50 Hz power in the dHPC corresponding to avoidance of a shock zone, suggesting that gamma can reflect behavioral output. Here, we show that gamma power increases after an incorrect choice. The direction and strength of power was predictive of accuracy on the next trial. This association indicates that gamma is involved in short-term, real-time processing needed when deciding the next choice. Contrasting with gamma, beta may be indicative of a neural state that facilitates memory processes.

## Conclusion

This study demonstrates that specific frequency bands in the LFPs of dHPC are selectively engaged during different behavioral time scales when rats perform a complex spatial set shifting task. Session-based alignment of LFPs to salient task milestones (e.g. the block switch or learning point) show that dHPC theta and beta power decrease before, with theta power also increasing after, an update in task context and reward contingency rules. Beta power in dHPC increases around the time rats learn the current rule, further corroborated by the observation of higher beta power during exploration trials. We also found an association between choice accuracy and the strength of dHPC gamma power during the preceding return, but not choice, epoch. Overall, our results support the hypothesis that specific dHPC rhythmic frequencies contribute differentially to distinct timescales of behavioral and task-relevant information, demonstrating the importance of understanding each frequency band’s role in memory across different behavioral timescales

## Acknowledgements

We would like to thank the following for their contributions to this work: Ginger Mullins, Sky Andrews, Kyra Diaz, and Jillian Perrone for their assistance in animal care and training; Kevan Kidder for his technical assistance; Victoria Hones for her support in animal care, training, and comments on the manuscript. This work was supported by the following: NIMH grant MH119391 to SJYM and the University of Washington Mary Gates Endowment for Students Undergraduate Research Travel Award to MB. There are no conflicts of interest to report.

## Author contributions

MB collected data, performed formal analyses, and wrote the original draft of the manuscript. JTM collected data and contributed revisions to the manuscript. SJYM supervised the project and contributed revisions to the manuscript.

